# Prediction and updating abilities in motor imagery during the Timed Up and Go task in young and older adults

**DOI:** 10.1101/2025.07.24.666513

**Authors:** Pierre-Olivier Morin, Christine Assaiante, Romain Brasselet, Angelo Arleo, Denis Sheynikhovich, Catherine Persephone Agathos

## Abstract

Aging is associated with declines in sensorimotor and cognitive functions that affect internal motor models, thought to mediate the risk of falls. Motor imagery, an experimental window into internal models, has been studied in aging, but it remains unclear whether age differentially impacts their predictive versus updating components. In this study, younger and older adults completed the Timed Up and Go task along with imagined trials before and after execution, enabling separate assessment of prediction and update accuracy. It was found that older adults exhibited similar or better accuracy during prediction and update, but the accuracy measures were linked with distinct cognitive and sensorimotor factors in the two age groups. These findings suggest that while internal model function is preserved in healthy aging, at least for every-day tasks, it is shaped by different compensatory mechanisms across the lifespan.

## Introduction

Aging is associated with sensory, motor, and cognitive changes that contribute to deficits in postural and locomotor control, thereby increasing the risk of falls in older adults (Gadkaree et al., 2016; Paraskevoudi et al., 2018; Seidler et al., 2010; Zapparoli et al., 2022). Falls represent a major public health concern, with 30 to 50% of individuals over the age of 65 experiencing at least one injurious fall each year (Stevens et al., 2008). These functional declines are thought to be linked to age-related changes in motor affordances and their control—for example, a shift from continuous to more intermittent motor control strategies (Boisgontier & Nougier, 2013) or a reliance on outdated motor representations that reflect previous, rather than current, capabilities (Caffier et al., 2019). Underlying these changes are sensorimotor representations, or internal models of action, which are essential for motor control by enabling individuals to predict, plan, and execute movements effectively (Miall & Wolpert, 1996; Wolpert et al., 1995). In older adults, these internal models may become less accurate or more resistant to updating, potentially resulting in over-optimistic predictions about future actions (Lafargue et al., 2013).

While a direct experimental assessment of age effects on internal models is difficult, they can be studied via motor imagery – i.e., mental simulation of movement without physical execution (Jeannerod, 1995) – a valuable paradigm for assessing internal representations of action. This process relies on a neural network that significantly overlaps with the circuitry involved in actual motor execution (Hardwick et al., 2018). Due to this shared neural architecture, motor imagery is thought to contribute to the construction and refinement of internal models, much like sensory feedback from executed movements (Shadmehr & Krakauer, 2008; Wolpert et al., 2011). Some experimental evidence suggests that internal models, as assessed by motor imagery, are perturbed or altered in the aging brain (Skoura et al., 2005, Boisgontier & Nougier, 2013), in accord with the evidence that impaired motor imagery performance is associated with higher fall risk (Grenier et al., 2018; Nakano et al., 2020; Sakurai et al., 2017). However, it is not clear whether internal model execution (i.e. model-based prediction) or updating (i.e. model recalibration based on sensory feedback) are affected (Vandevoorde & Orban de Xivry, 2019). Moreover, motor imagery depends on visuo-cognitive and sensorimotor integration processes, affected by aging to a large degree (Costello & Bloesch, 2017; Ren et al., 2013), but the role of these processes in execution and updating of internal models is not clear. Indeed, cognitive decline in older age has been linked with deficits in motor imagery (Allali et al., 2012; Beauchet et al., 2014; Lallart et al., 2012), associated with deteriorated working memory of mental representations of movements and mental flexibility (Schott, 2012, Malouin et al., 2010, Rudiger et al., 2017). In addition, age-related changes in proprioception and sensorimotor integration were shown to lead to motor control deficits, potentially via their contribution to the construction and updating of internal representations of action (Goble et al., 2009, Seidler et al., 2010).

Motor imagery is typically examined along the dimensions of vividness of motor representation, controllability, timing and temporal congruence (McAvinue & Robertson, 2008; Saimpont et al., 2015). The temporal congruence between motor imagery and motor execution is particularly relevant to fall risk as it reflects subject’s abilities in motor planning and adaptation, central to balance maintenance and safe locomotion. However, testing the effect of age on temporal congruence produced inconsistent results, with some studies reporting that older adults overestimate motor task duration compared to young adults (Personnier et al., 2010), while others observe an underestimation of execution time in older adults, especially in those over 80 years of age (Sakurai et al., 2017; Schott, 2012). Yet other studies report an equivalent performance between age groups (Malouin & Richards, 2010; Skoura et al., 2005). The reasons for these discrepancies are not clear and they may in fact be caused by differences in cognitive and sensorimotor profiles of the participants.

To evaluate the effect of aging on execution and updating of internal models, as well as to assess the contribution of cognitive and sensorimotor factors to these processes, our study investigated the temporal congruence between imagined and executed Timed Up and Go (TUG) task in young and older adults. This test is a common functional balance assessment due to its simplicity and relevance in representing daily living activities and postural challenges in older age (Podsiadlo & Richardson, 1991). Accordingly, performance in this test is a good predictor of fall risk (Beauchet et al., 2011; Beck Jepsen et al., 2022; Montero-Odasso et al., 2022). In contrast to previous studies (Personnier et al., 2008; Saimpont et al., 2013; Skoura et al., 2005), we assessed sequential prediction and updating of internal motor representations by examining individuals’ performance when imagery was conducted before and after motor task execution. This approach allowed us to examine whether age-related deficits in motor imagery are primarily due to impaired prediction, compromised updating, or both. Furthermore, by characterizing the cognitive and sensorimotor profiles of participants, we sought to identify which factors—such as working memory, executive function, proprioception, and visuo-spatial processing—most strongly influence internal model function in older adults. Understanding these mechanisms may offer valuable insights into fall risk and inform the development of more targeted interventions to preserve motor planning and control in aging populations.

## Methods

### Participants

Fifty-one older adults (77.78 ± 4.2 years, range 66-86 years, 28 females), and thirty-four young adults (30.29 ± 5.8 years, range 22-42 years, 19 females), participated in this study. The participants were part of the SilverSight cohort study conducted at the Vision Institute and Quinze-Vingts National Ophthalmology Hospital in Paris, France, started in 2015 (Lagrené et al., 2019). All participants had normal or corrected-to-normal vision and had no history of sensory, motor, neurological or psychiatric dysfunctions. All participants included in this study scored 24 or higher on the Mini Mental State Examination (MMSE), indicating a generally intact cognitive function.

The following visuo-cognitive measures were evaluated for each participant: spatial working memory (forward and backward span on the Corsi block-tapping test, Corsi, 1972), inhibition capacity (Go/No go task, Diamond, 2013), concerns about falling (Falls Efficacy Scale - International, Greenberg, 2012), propensity for conscious control and monitoring of movements (Movement Self-Consciousness and Conscious Motor Processing Scales of the Movement-Specific Reinvestment questionnaire, Masters et al., 2005) and motor imagery capacity (Kinesthetic, and 1st and 3rd person Visual scales of the Motor Imagery Questionnaire, Robin et al., 2021). Proprioceptive ability was assessed by plantar sensitivity at the first metatarsal head (MT1) using an epicritic tactile sensation test (Snyder et al., 2016). Walking speed was measured during free walking over 9m distance. Performance on these tests and their comparison across age-groups, together with demographic profiles of participants are provided in Table 1. All experimental procedures were conducted in accordance with the tenets of the Declaration of Helsinki and approved by the Ethical Committee “CPP Ile de France V” (ID_RCB 2015-A01094-45, CPP N: 16122). All volunteers provided informed, written consent to participate.

**Table 1.**
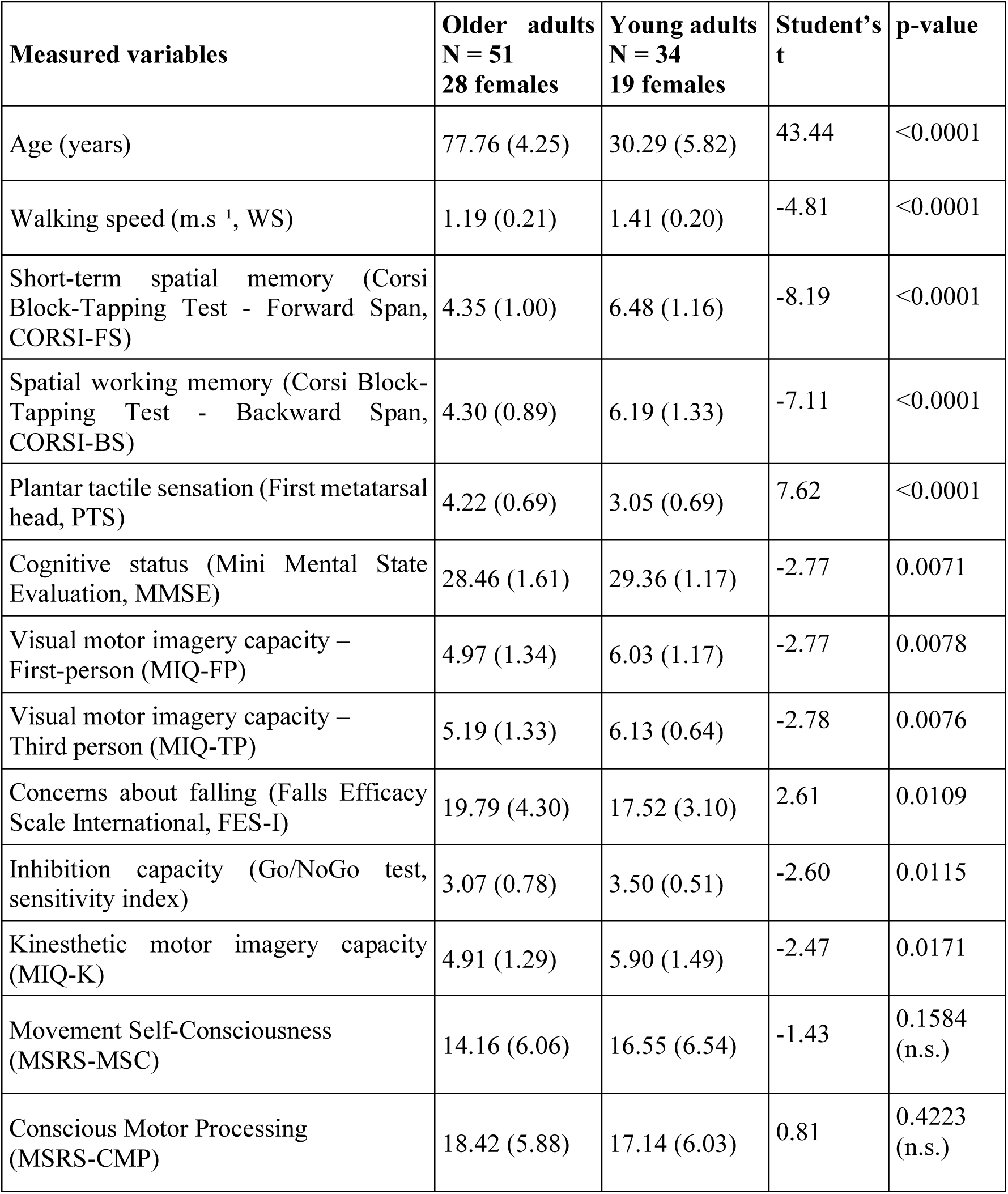
Cognitive and sensorimotor variables measured for each participant. Data are presented as means with standard deviations in parenthesis. Higher assessment score values represent better performance except for: plantar tactile sensation (higher values correspond to reduced sensitivity) and falls efficacy scale (higher values indicate greater concerns about falling). The variables are presented in the order of decreasing significance.

There were missing data due to participant availability constraints. Specifically, the Movement-Specific Reinvestment questionnaire data was missing in 13 (25%) older and 12 (35%) younger adults; the Motor Imagery questionnaire was missing for 16 (31%) older and 17 (50%) younger participants. Additionally, MMSE and Falls Efficacy Scale assessments were missing in 3 older and 1 younger participant, respectively. Missing data were handled using pairwise deletion for statistical analyses.

### Experimental design and procedure

The protocol for the TUG test, as described by Podsiadlo & Richardson (1991), was explained to the participants: rise from a seated position, walk 3 m, turn around, walk back, and sit down. In order to test individuals’ motor imagery in terms of predictive ability, the experimenter demonstrated the task instead of having participants perform the familiarization trial outlined in the original protocol. Participants were informed that they would perform three repetitions of the TUG by physically moving and four imagined repetitions (iTUG) – before and after each executed TUG (see Figure 1). In both cases, participants were given a “go” signal and were asked to say “stop” at the end of the trial when their back touched the seat’s backrest (executed condition) or when they imagined this happening (Beauchet et al., 2010). The iTUGs were performed while seated in the same starting position as the TUG, with eyes closed. For the imagined versions, participants were instructed to imagine how they *would perform* (on the first iTUG trial, prediction) or how they *had just performed* (after each executed trial, update) the TUG. The experimenter initiated the stopwatch right after the “go” signal and stopped it when the participant verbally indicated “stop”. Participants were given no feedback on their performance.

**Figure 1:**
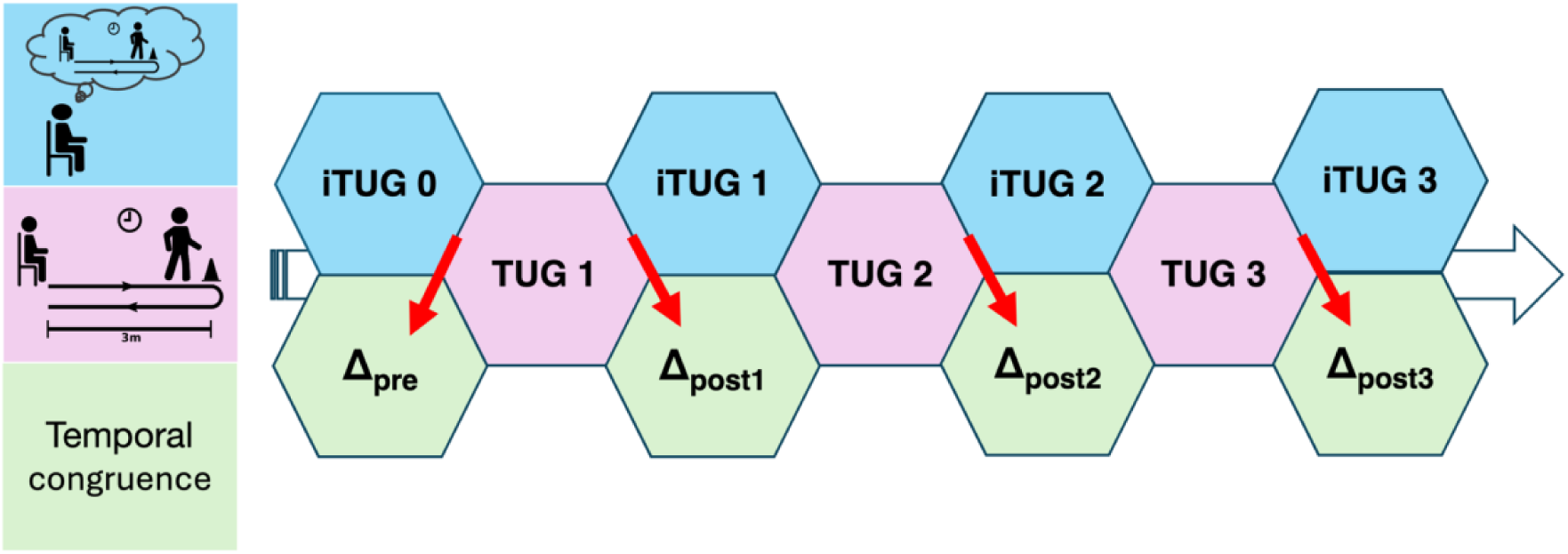
A schematic representation of the experimental procedure. First, participants are asked to imagine the TUG task (iTUG 0: predicted duration), followed by 3 repetitions of physically executed (TUG1 – TUG3: actual duration) and imagined (iTUG1 – iTUG3: updated duration) tasks. The temporal congruence of prediction is evaluated based on the difference between the first actual and predicted durations (Δ_pre_). The temporal congruence of update is evaluated based on the difference between the actual and updated durations (Δ_post1_, Δ_post2_, Δ_post3_)

### Variables of interest

The variables of interest included the duration of each TUG and iTUG trial, recorded using a stopwatch with 0.01s precision. Additionally, weighted differences between TUG and iTUG durations were calculated to define the temporal congruence between the imagined and executed tasks. To assess participants’ predictive motor imagery ability, a time difference was calculated between the first imagined and the first executed version of the TUG (henceforth termed *prediction error*), according to the equation (Beauchet et al., 2010):

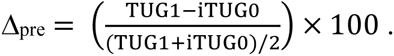

For subsequent trials, the time difference determining participants’ ability to update their movement representation were calculated between each executed and subsequent imagined trial (henceforth termed *update error*), according to the following equation:

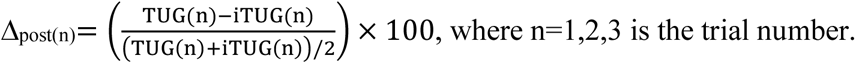

### Data analysis

All statistical analyses were performed using R (R Core Team, version 4.5.0). Mixed ANOVAs were performed using *rstatix* package with age group as a between-subjects factor and trial as a within-subjects factor, with paired comparisons implemented using Holm-Bonferroni procedure. Effect sizes are reported using partial eta squared (ηp²). For correlation analyses, Pearson’s or Spearman’s tests were performed depending on the data normality assumptions. Correlation matrices were compared using Mantel permutation test (*vegan* package version 2.6-10) with Spearman’s ⍴ as the correlation measure. The statistical difference between correlation coefficients was assessed using *cocor* package (version 1.1-4). For the comparisons between age groups (independent), Fisher’s z is reported, while for the comparisons between prediction and update errors within a group (dependent overlapping), Hittner’s z is reported.

Since no significant sex-related effects emerged across our analyses, sex was not retained as a factor in the final models presented here.

## Results

### Executed and imagined TUG duration

Participants’ performance on the executed TUG task is shown in Figure 2A. Consistent with general age-related slowing while walking (Agathos et al., 2023) we observed a significant main effect of age (F(1, 86) = 27.95, p < .001; ηp² = 0.245), with older adults exhibiting longer TUG durations compared to young adults (older adult μ ± sd: 11.30 ± 2.20 s; young adult μ ± sd: 9.14 ± 1.72 s). A significant main effect of trial number was also observed (F(2, 172) = 17.60, p < .001; ηp² = 0.17). Paired comparisons revealed that TUG durations were significantly different across all three trials, with TUG duration reducing over time likely due to a habituation effect (TUG1: 10.8 ± 2.38 s; TUG2: 10.4 ± 2.20 s; TUG3: 10.0 ± 2.18 s). This effect was statistically equivalent in both age groups, as the interaction between age group and trial number was not significant (F(2, 172) = 2.25, p = 0.11).

**Figure 2:**
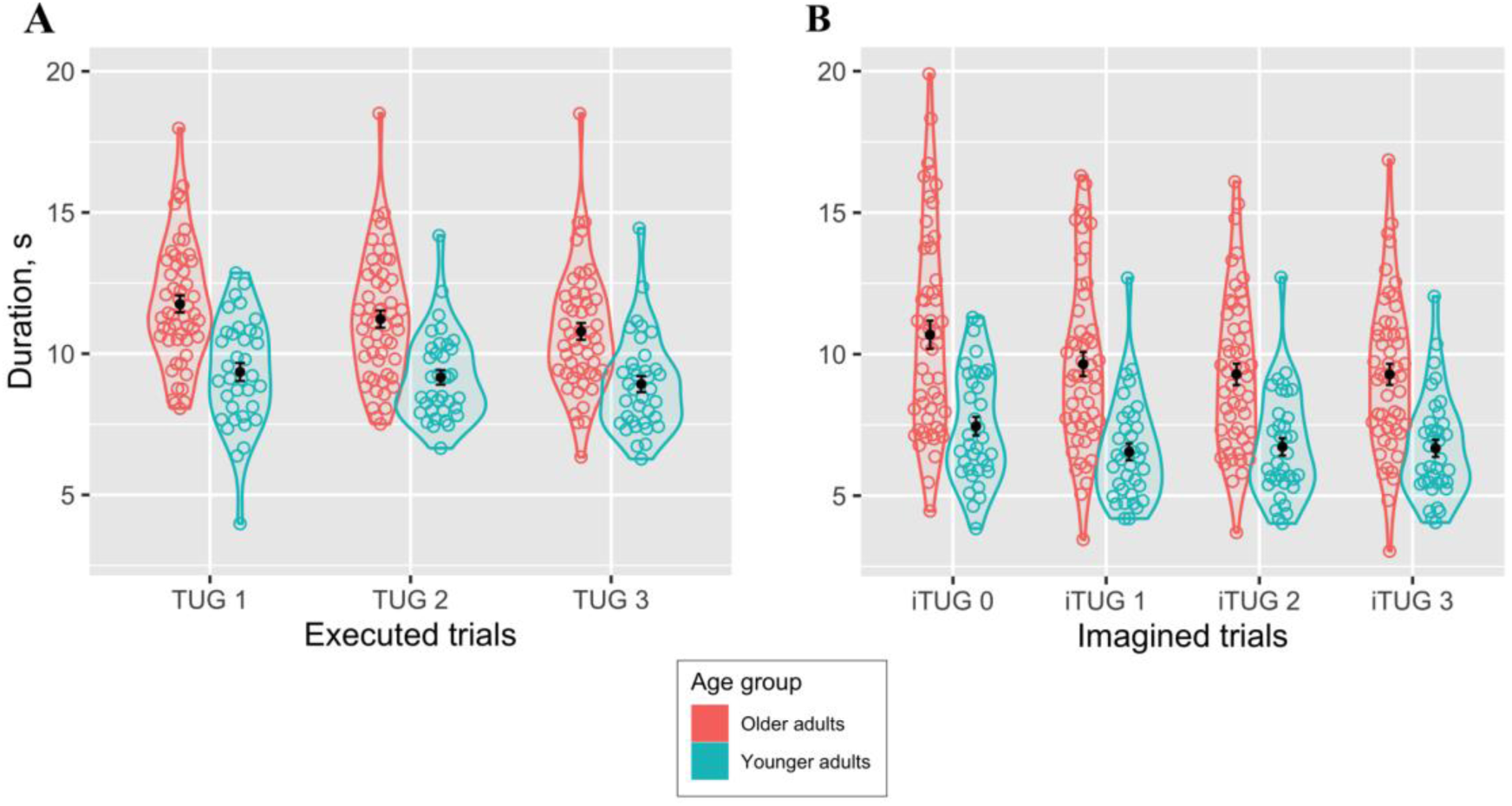
Executed (**A**) and imagined (**B**) TUG duration (in seconds) as a function of trial for the two age groups. Small circles correspond to different subjects, mean (±SE) are shown in black.

For the imagined TUG task (Fig. 2B), participants were first asked to imagine performing the TUG, whereas in the subsequent three trials they were asked to imagine how they had just executed the TUG. When comparing iTUG durations across the two age groups and four trials, we found a main effect of age group on iTUG duration (F(1, 86) = 31.12, p < .001; ηp² = 0.27), whereby older adults exhibited significantly longer iTUG durations compared to young adults (older adult: 9.73 ± 3.07 s; young adult: 6.85 ± 1.88 s). We also observed a significant main effect of trial number (F(3, 258) = 19.22, p < .001; ηp² = 0.18). Post hoc comparisons revealed that, on average, the duration of the iTUG was longer on the first, prediction trial, compared to the three following update trials (iTUG1: 9.36 ± 3.41 s; iTUG2: 8.38 ± 3.02 s; iTUG3: 8.24 ± 2.71 s; iTUG4: 8.22 ± 2.70 s). The interaction between age group and trial number was not statistically significant (F(3, 258) = 0.86, p = .46). These results suggest that in both groups, internal models consistently reflected the real difference in TUG duration and that there was a strong effect of the first executed TUG, but not the subsequent ones, on the internal model.

In order to get an insight into the individual relationships between executed and predicted TUG duration in the two age groups, linear regression analyses were performed. When comparing the imagined with the first executed TUG (prediction performance, Fig. 3A), we found that correlation coefficients were significantly different between age groups (Fisher’s z = 2.399, p = 0.016). Moreover, the regression slope was significantly smaller than unity only in the younger group (t-ratio -3.631, p= 0.0009), suggesting that older adults are in fact better at predicting their motor performance. Similar results were obtained when comparing the executed TUG with the following imagined TUG (update performance, Fig. 3B, Fisher’s z = 2.3443, p = 0.019; t-ratio = -3.519, p= 0.001). These results show that in the older group, the longer a participant’s executed TUG was, the longer were his or her imagined TUG durations and by approximately the same amount of time for both prediction and update. In contrast, in the younger participants, imagined TUG durations were shorter compared to executed durations, as classically reported in the motor imagery literature including locomotion and other tasks (Guillot & Collet, 2005; Marchetti et al., 2022)

**Figure 3:**
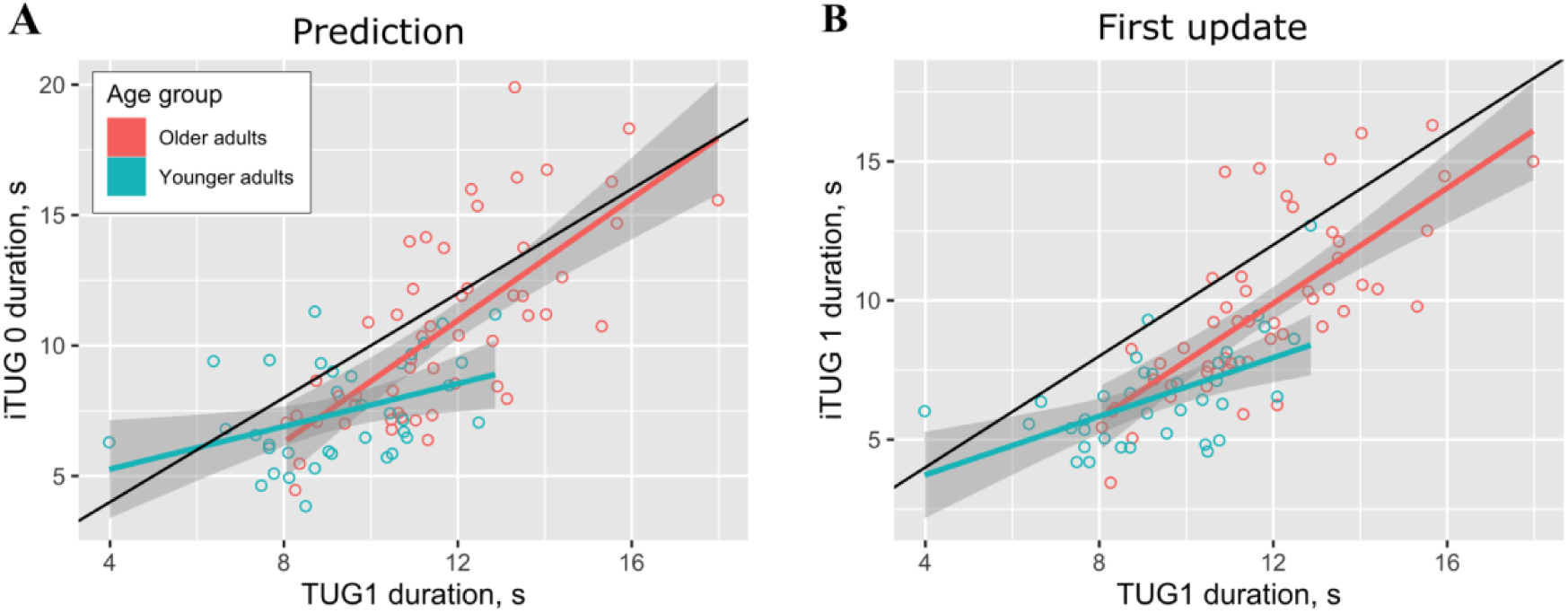
**A.** Predicted (imagined) TUG duration (iTUG 0) as a function of the actual duration in the first trial (TUG 1). **B.** Updated (imagined) TUG duration (iTUG 1) as a function of the same variable as in A. Each small circle corresponds to a participant from the older (in red) or the younger (in blue) group. The black line corresponds to the one-to-one relationship, corresponding to perfect prediction and update abilities.

### Temporal congruence

Temporal congruence, defined as the difference between executed and imagined durations (Saimpont et al., 2013), is a more direct measure of internal model accuracy. On average, participants of both age groups underestimated the duration of the TUG task when imagining it, as shown by the positive mean differences in Figure 4A. In the subsequent analysis, we separated the temporal congruence of prediction and update corresponding to conceptually distinct variables. While the average prediction error (Δ_pre_) was smaller in the older participants compared to younger ones (older adults: 13.2 ± 23.2; young adults: 23.3 ± 27.9), this difference did not reach statistical significance (t=-1.85, p=0.066).

**Figure 4:**
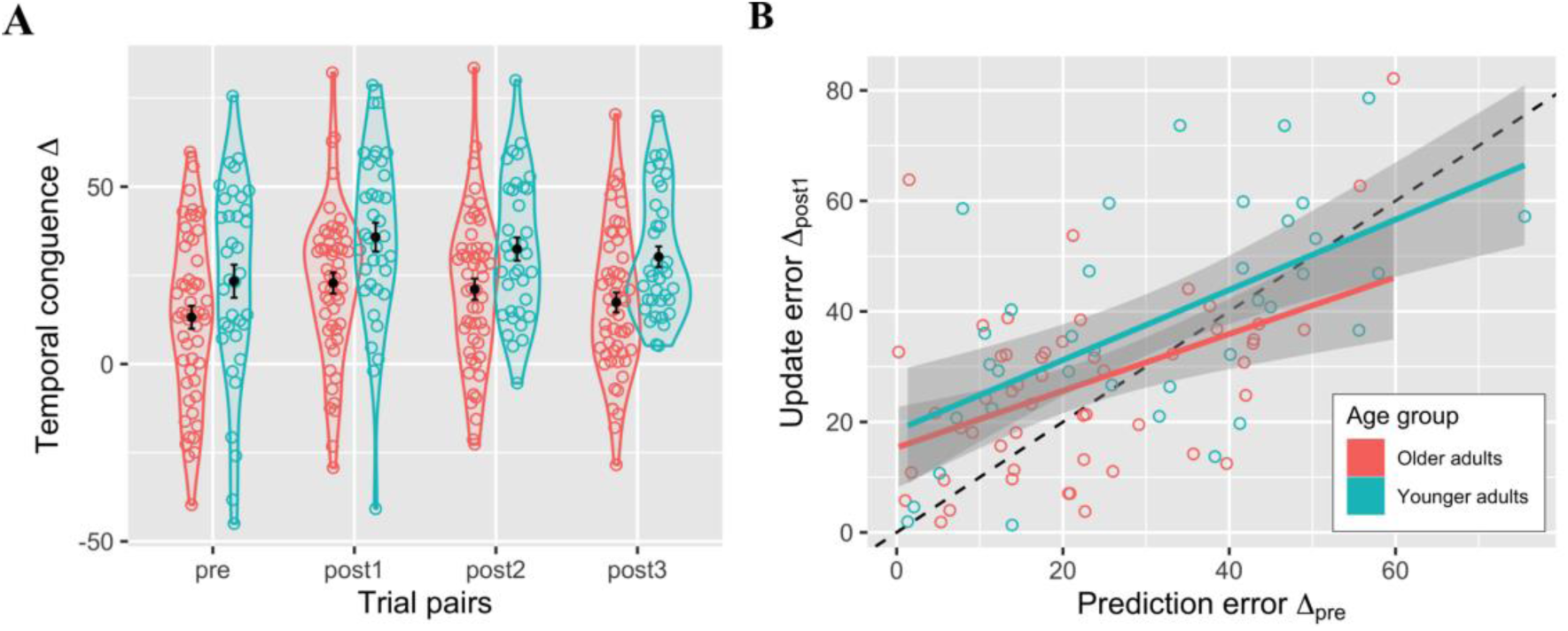
**A.** Temporal congruence between executed and imagined TUG durations, corresponding to the prediction trial pair (Δ_pre_) and three update pairs (Δ_post1_, Δ_post2_, Δ_post3_), see Methods. Mean (±SE) are shown in black. **B.** Update error (Δ_post1_) as a function of the prediction error (Δ_post1_). Each small circle corresponds to a participant from the older (in red) or the younger (in blue) group. The dotted black line corresponds to the one-to-one relationship, corresponding to identical prediction and update errors.

Regarding the update error (Δ_post_), we found a significant main effect of age group (F(1, 86) = 9.48, p = 0.003, ηp² = 0.10), with older adults exhibiting a smaller error compared to young adults (older adults: 20.4 ± 21.1; young adults: 32.8 ± 20.5). There was also a significant main effect of trial number (F(2, 172) = 4.83, p < 0.009, ηp² = 0.05), but no interaction (F(2,172)=0.13; p=0.88). Post-hoc comparisons indicated a significant difference between Δpost1 and Δ_post3_ (t(174) = 2.87, p = 0.015), revealing an improvement in temporal congruence across update trials (Δ_post1_ = 28.1 ± 23.4; Δ_post2_ = 25.7 ± 21.4; Δ_post3_ = 22.6 ± 20.1) in both age groups.

Finally, we looked at the relationship between the absolute values of temporal congruence of prediction and update at the individual level (Fig. 4B). In both age groups there was a highly significant correlation between Δ_pre_ and Δ_post1_ (older: r=0.77, p<10^-7^, younger: r=0.70, p<10^-5^) but no difference between the correlation coefficients across groups (Fisher’s z = -0.216, p = 0.83). Overall, these results reinforce the conclusion from the previous section that older adults possess a better internal model in relation to the TUG task (similar and smaller prediction and update errors, respectively, compared to younger adults) while exhibiting similar improvement over time and similar individual ratios for prediction and update in both ages.

### Explaining temporal congruence with cognitive and sensorimotor variables

The results above suggest that, at least for the TUG task, motor imagery is more accurate in the older than in the younger group in terms of temporal congruence. However, as shown in Table 1, most of the variables measuring cognitive, sensorimotor and psychological functions, potentially related to the TUG task, differ strongly between the two age groups. An interesting question is whether the apparently age-resistant performance in motor imagery prediction and update relies on compensatory changes in (a subset of) these variables.

To address this question, we computed correlation matrices of variables in Table 1 separately in the two age groups, with the addition of the absolute values of Δ_pre_ and Δ_post1_ as new columns, and evaluated whether the overall correlation structure differed across the groups. The test was not significant (Mantel’s r=0.10, p=0.136), suggesting that the null hypothesis of there being no statistical relationship between the two correlation matrices cannot be rejected. While this does not mean the two matrices are not related in general, the current data suggest that age may have a substantial effect on the overall correlation structure of the cognitive, sensorimotor variables, prediction and update measures.

Our subsequent analysis focused on the correlation of each of the 13 variables in Table 1 with the prediction and update errors in the two age groups. More specifically, for each variable separately, we assessed the correlation *(i)* between the variable and the prediction error, as well as *(ii)* between the variable and the update error. We also checked whether *(iii)* the influence of the variable on prediction is statistically different from its influence on update errors. The results of these comparisons are reported in Table 2. It can be seen that in older participants, prediction errors are associated with greater age, while update errors tend to be linked with visual motor imagery (FP). In contrast, no such associations were observed in the younger group. Instead, there was a statistical tendency for the correlation of prediction errors with the proprioceptive sensitivity, while update errors were associated with the movement self-consciousness score.

**Table 2.** Linear correlation coefficients for the dependencies between the sensorimotor variables (first column) and the prediction error (**Δ_pre_**) as well as the update error (**Δ_post1_**). Pearson or Spearman correlation is reported depending on the data normality. The column **Δ_pre_ vs Δ_post1_** reports whether the correlation coefficients in the first two columns are statistically different (Hittner test statistic is reported). Only correlations with p<0.1 are reported.

| Variable | Younger adults |  |  | Older adults |  |  |
| --- | --- | --- | --- | --- | --- | --- |
| | $\Delta_{pre}$ | $\Delta_{post}$ | $\Delta_{pre}$ vs $\Delta_{post}$ | $\Delta_{pre}$ | $\Delta_{post}$ | $\Delta_{pre}$ vs $\Delta_{post}$ |
| Age | n. s. | n. s. | n. s. | $\rho=0.34$<br>$p=0.015$ | n. s. | $z=2.148$<br>$p=0.031$ |
| MSRS-MSC | n. s. | $\rho=-0.43$<br>$p=0.037$ | $z=2.128$<br>$p=0.033$ | n. s. | n. s. | - |
| MIQ-FP | n. s. | n. s. | n. s. | n. s. | $\rho=0.28$<br>$p=0.097$ | $z=-2.189$<br>$p=0.029$ |
| MIQ-TP | n. s. | n. s. | n. s. | n. s. | n. s. | - |
| MIQ-K | n. s. | n. s. | n. s. | n. s. | n. s. | - |
| PTS | $r=-0.28$<br>$p=0.072$ | n. s. | $z=-1.87$<br>$p=0.061$ | n. s. | n. s. | n. s. |

## Discussion

This study examined motor imagery ability in terms of timing and temporal conguence using the Timed Up and Go test in young and older adults. We sought to understand age-related differences as well as their cognitive and sensorimotor correlates for the prediction and updating of motor representations. Our findings reveal differences in motor imagery between age groups, both before and after executing the TUG task, with the older population exhibiting an overall more accurate internal model of action. Participants generally underestimated the actual duration of the TUG while imagining it, in accordance with previous evidence on motor imagery (Guillot & Collet, 2005). Contrary to expectations, however, older adults exhibited better temporal congruence than younger adults, as their updated estimates were closer to actual TUG duration on average. Moreover, on the individual level, older adults more accurately imagined the TUG task both before and after its execution. Finally, age-related differences in temporal congruence were associated with specific factors such as movement self-consciousness, visual imagery and proprioception, but not with other cognitive and sensorimotor variables.

### More accurate internal model and its recalibration in older adults

Older adults demonstrated a more accurate internal representation of action, as measured by temporal congruence, than young adults in both prediction and updating of motor imagery. These findings support and extend previous data suggesting similar (Schott & Munzert, 2007) or superior (Watanabe & Tani, 2022) motor imagery abilities in older adults and challenges traditional views of age-related decline in motor-cognitive functions (Zhuang et al., 2020). It is likely that lifetime experience may lead to more robust temporal motor representations when it comes to basic functional tests such those represented by the TUG task. This aligns with previous literature suggesting that older adults maintain good temporal congruence during familiar, habitual movements. For example, older adults exhibited reduced temporal congruence (*i.e.,* larger differences) between imagined and executed movements only when these are constrained, but not during free/habitual movements (Saimpont et al., 2013). The TUG task used in our study represents a familiar, everyday movement sequence, which may explain smaller temporal errors observed in the older participants. This experience-based advantage may reflect compensatory mechanisms in aging, whereby extensive practice with specific motor tasks can preserve performance despite general age-related decline (Krampe & Ericsson, 1996).

Analysis of prediction and update accuracy show that when the duration of the executed TUG increased, the relationship between the actual and imagined durations approached an asymptote with a slope of 1, reflecting near-perfect temporal congruence. Accordingly, correlations between the first executed TUG and the first (prediction) as well as second (update) imagined durations revealed that, in older adults, slower actual performance was associated with a better accuracy in imagined durations. This finding suggests that individuals who face a greater difficulty during real execution may produce more elaborate and detailed motor simulations. Having fewer reliable sensory inputs and reduced ability to adjust movements through feedback, these individuals might rely more heavily on their motor representations to compensate. In this view, more accurate imagery reflects not only caution but also the necessity to pre-emptively plan actions in a more exhaustive manner. This suggests that cognitively intact older adults like those in our study may favor anticipatory rather than feed-back strategy, on the basis of preserved internal representations, to optimize their motor control.

### Factors contributing to age-related changes in temporal congruence

Age itself emerged as an important factor in older adults, but only for prediction error, with no influence on updating. This aligns with previous literature suggesting that temporal congruence deteriorates significantly beyond certain age thresholds, potentially around 80 years (Sakurai et al., 2017; Schott, 2012). The absence of correlation between age and update errors suggests that while initial prediction abilities may decline with advancing age, the updating process itself remains relatively preserved. This is further supported by the lack of interaction effect between age group and trial number on updating errors.

When examining the role of motor imagery vividness, no significant correlations were observed between MIQ scores and prediction or update errors in either age group. However, the *difference* between the associations of vividness scores with prediction and update errors – particularly for the MIQ-FP subscale – was statistically significant in the older group, suggesting that the influence of vividness may be more pronounced during the updating phase of motor imagery than during prospective estimation. These findings support the view that vividness of motor imagery, while often considered beneficial for engaging motor-related brain networks and performance in sport and rehabilitation contexts (Guillot & Collet, 2005), does not necessarily enhance temporal accuracy, particularly in older adults. Prior work has shown that imagery vividness and temporal congruence constitute distinct but complementary dimensions of motor imagery, underpinned by partially dissociable neural mechanisms (Subirats et al., 2018; Zabicki et al., 2019). As highlighted by Williams et al., (2015), vividness and temporal congruence offer complementary rather than redundant insights into imagery quality, and our findings align with this distinction.

Movement self-consciousness contributed to better motor imagery in the younger, but not in the older, group. The inclusion of the MSRS in our study aimed to capture individuals’ tendencies to consciously monitor or think about their own movements, which may influence motor imagery processes. The Movement Self-Consciousness (MSC) subscale captures the degree to which individuals are concerned with how their movements appear to others. The negative correlation with update errors suggests that a greater insecurity about one’s movements facilitates more accurate updating of motor representations following execution. Interestingly, this factor emerged only in relation with the updating error – after the TUG was executed and thus created a more conscious consideration of one’s movements.

Finally, the proprioceptive ability, measured by plantar sensitivity, emerged as a potentially important factor, in accord with the importance of somatosensory signals for internal models of action (Cardinali et al., 2016). In younger adults, and in relation to prediction but not updating, greater tactile sensitivity (lower threshold) was associated with less accurate representations of movement duration. As tactile sensitivity is a measure of somatosensory perception, a contribution to better temporal congruence in motor imagery would be expected. Our finding could thus reflect a compensatory anticipatory strategy: individuals with reduced plantar sensitivity utilize prospective planning, rather than the proprioceptive feedback. Conversely, individuals with good tactile sensitivity may rely more on feedback-based correction. Reduced tactile perception at the foot, particularly at the first metatarsal head (MT1), is strongly associated with balance deficits and slower walking speed, both of which contribute to increased fall risk (Cruz-Almeida et al., 2014). The lack of effect in our study may be caused by higher sensitivity thresholds in older adults, thus not contributing to the imagery process. Alternatively, older adults may have developed compensatory strategies that rely less on immediate sensory feedback and more on established motor representations (Vandevoorde & De Xivry, 2020). Yet another interpretation is related to age-related changes in multisensory integration processes. Recent developmental studies highlight a long maturational process of the cortical correlates underlying sensory integration and motor anticipation (Cignetti et al., 2018; Fortin et al., 2021) and the development of internal representations (Cignetti et al., 2017; Fontan et al., 2017) from late childhood to adulthood, with a protracted development of the proprioceptive brain network during and beyond adolescence. Advanced age can thus lead to multisensory reweighting, minimizing the effect of unreliable tactile information in the construction and updating of internal models.

### Study limitations and future directions

Several limitations of the present study should be acknowledged. Participants in our older adult sample were screened for the absence of motor, neurophysiological and ophthalmological deficits (Lagrené et al., 2019), potentially limiting generalizability to frailer older populations in which motor imagery deficits may be more pronounced. Additionally, the TUG task, while ecologically valid, represents only one type of movement. Future studies are required to examine temporal congruence during prediction and update across motor tasks with increased behavioral constraints (Saimpont et al., 2013). Examining neural mechanisms underlying preserved motor imagery while accounting for individual phenotypes can further aid in the design of motor imagery training as a therapeutic intervention. Finally, longitudinal studies tracking changes in temporal congruence alongside cognitive and sensorimotor measures would provide valuable insights into the developmental trajectory of motor imagery abilities.

## Conclusion

This study provides novel evidence that older adults, despite general age-related declines in motor and cognitive function, retain— and in some cases exceed— the ability of younger adults to accurately predict and update internal models of movement, at least for well-learned tasks such as the Timed Up and Go. Contrary to expectations, older participants demonstrated greater temporal congruence between imagined and executed movements than younger participants, suggesting a potential compensatory role of motor planning mechanisms when sensory feedback becomes less reliable. Importantly, age-related changes in motor imagery performance were not uniformly associated with general cognitive or sensorimotor decline. Instead, select factors – such as movement self-consciousness, visual imagery ability, and proprioceptive sensitivity – emerged as key correlates, underscoring the importance of individual profiles in shaping internal model accuracy. These results suggest that older adults may shift toward anticipatory control strategies supported by robust motor representations, particularly in familiar, functional movements. Future work should explore how these preserved or enhanced imagery abilities might be leveraged in clinical interventions aimed at promoting safe mobility and functional independence in older adults.

## References

Agathos, C. P., Velisar, A., & Shanidze, N. M. (2023). A Comparison of Walking Behavior during the Instrumented TUG and Habitual Gait. Sensors, 23(16), 7261. 10.3390/s23167261

Allali, G., Laidet, M., Assal, F., Beauchet, O., Chofflon, M., Armand, S., & Lalive, P. H. (2012). Adapted Timed Up and Go: A Rapid Clinical Test to Assess Gait and Cognition in Multiple Sclerosis. European Neurology, 67(2), 116–120. 10.1159/000334394

Beauchet, O., Annweiler, C., Assal, F., Bridenbaugh, S., Herrmann, F. R., Kressig, R. W., & Allali, G. (2010). Imagined Timed Up & Go test: A new tool to assess higher-level gait and balance disorders in older adults? Journal of the Neurological Sciences, 294(1–2), 102–106. 10.1016/j.jns.2010.03.021

Beauchet, O., Fantino, B., Allali, G., Muir, S. W., Montero-Odasso, M., & Annweiler, C. (2011). Timed up and go test and risk of falls in older adults: A systematic review. *Journal of Nutrition*, Health and Aging, 15(10), 933–938. 10.1007/s12603-011-0062-0

Beauchet, O., Launay, C. P., Sejdić, E., Allali, G., & Annweiler, C. (2014). Motor imagery of gait: A new way to detect mild cognitive impairment? Journal of NeuroEngineering and Rehabilitation, 11(1), 66. 10.1186/1743-0003-11-66

Beck Jepsen, D., Robinson, K., Ogliari, G., Montero-Odasso, M., Kamkar, N., Ryg, J., Freiberger, E., & Masud, T. (2022). Predicting falls in older adults: An umbrella review of instruments assessing gait, balance, and functional mobility. BMC Geriatrics, 22(1), 615. 10.1186/s12877-022-03271-5

Boisgontier, M. P., & Nougier, V. (2013). Ageing of internal models: From a continuous to an intermittent proprioceptive control of movement. AGE, 35(4), 1339–1355. 10.1007/s11357-012-9436-4

Caffier, D., Luyat, M., Crémoux, S., Gillet, C., Ido, G., Barbier, F., & Naveteur, J. (2019). Do Older People Accurately Estimate the Length of Their First Step during Gait Initiation? Experimental Aging Research, 45(4), 357–371. 10.1080/0361073X.2019.1627495

Cardinali, L., Brozzoli, C., Luauté, J., Roy, A. C., & Farnè, A. (2016). Proprioception Is Necessary for Body Schema Plasticity: Evidence from a Deafferented Patient. Frontiers in Human Neuroscience, 10. 10.3389/fnhum.2016.00272

Cignetti, F., Fontan, A., Menant, J., Nazarian, B., Anton, J.-L., Vaugoyeau, M., & Assaiante, C. (2017). Protracted Development of the Proprioceptive Brain Network During and Beyond Adolescence. Cerebral Cortex, 27(2), 1285–1296. 10.1093/cercor/bhv323

Cignetti, F., Vaugoyeau, M., Decker, L. M., Grosbras, M.-H., Girard, N., Chaix, Y., Péran, P., & Assaiante, C. (2018). Brain network connectivity associated with anticipatory postural control in children and adults. Cortex, 108, 210–221. 10.1016/j.cortex.2018.08.013

Corsi, P. M. (1972). Short Title by HUMAN MEMORY AND THE MEDIAL TEMPORAL REGION OF THE BPAIN [McGill University]. In Psychology. https://escholarship.mcgill.ca/concern/theses/05741s554

Costello, M. C., & Bloesch, E. K. (2017). Are older adults less embodied? A review of age effects through the lens of embodied cognition. Frontiers in Psychology, 8. 10.3389/fpsyg.2017.00267

Cruz-Almeida, Y., Black, M. L., Christou, E. A., & Clark, D. J. (2014). Site-specific differences in the association between plantar tactile perception and mobility function in older adults. Frontiers in Aging Neuroscience, 6. 10.3389/fnagi.2014.00068

Diamond, A. (2013). Executive Functions. Annual Review of Psychology, 64(Volume 64, 2013), 135–168. 10.1146/annurev-psych-113011-143750

Fontan, A., Cignetti, F., Nazarian, B., Anton, J.-L., Vaugoyeau, M., & Assaiante, C. (2017). How does the body representation system develop in the human brain? Developmental Cognitive Neuroscience, 24, 118–128. 10.1016/j.dcn.2017.02.010

Fortin, C., Barlaam, F., Vaugoyeau, M., & Assaiante, C. (2021). Neurodevelopment of Posture-movement Coordination from Late Childhood to Adulthood as Assessed From Bimanual Load-lifting Task: An Event-related Potential Study. Neuroscience, 457, 125–138. 10.1016/j.neuroscience.2020.12.030

Gadkaree, S. K., Sun, D. Q., Li, C., Lin, F. R., Ferrucci, L., Simonsick, E. M., & Agrawal, Y. (2016). Does Sensory Function Decline Independently or Concomitantly with Age? Data from the Baltimore Longitudinal Study of Aging. Journal of Aging Research, 2016, e1865038. 10.1155/2016/1865038

Goble, D. J., Coxon, J. P., Wenderoth, N., Van Impe, A., & Swinnen, S. P. (2009). Proprioceptive sensibility in the elderly: Degeneration, functional consequences and plastic-adaptive processes. Neuroscience and Biobehavioral Reviews, 33(3), 271–278. 10.1016/j.neubiorev.2008.08.012

Greenberg, S. A. (2012). Analysis of Measurement Tools of Fear of Falling for High-Risk, Community-Dwelling Older Adults. Clinical Nursing Research, 21(1), 113–130. 10.1177/1054773811433824

Grenier, S., Richard-Devantoy, S., Nadeau, A., Payette, M.-C., Benyebdri, F., Duhaime, M.-M. B., Gunther, B., & Beauchet, O. (2018). The association between fear of falling and motor imagery abilities in older community-dwelling individuals. Maturitas, 110, 18–20. 10.1016/j.maturitas.2018.01.001

Guillot, A., & Collet, C. (2005). Duration of mentally simulated movement: A review. Journal of Motor Behavior, 37(1), 10–20. 10.3200/JMBR.37.1.10-20

Hardwick, R. M., Caspers, S., Eickhoff, S. B., & Swinnen, S. P. (2018). Neural correlates of action: Comparing meta-analyses of imagery, observation, and execution. Neuroscience and Biobehavioral Reviews, 94, 31–44. 10.1016/j.neubiorev.2018.08.003

Jeannerod, M. (1995). Mental imagery in the motor context. Neuropsychologia, 33(11), 1419–1432. 10.1016/0028-3932(95)00073-C

Krampe, R. T., & Ericsson, K. A. (1996). Maintaining excellence: Deliberate practice and elite performance in young and older pianists. Journal of Experimental Psychology. General, 125(4), 331–359. 10.1037//0096-3445.125.4.331

Lafargue, G., Noël, M., & Luyat, M. (2013). In the Elderly, Failure to Update Internal Models Leads to Over-Optimistic Predictions about Upcoming Actions. PLoS ONE, 8(1), e51218. 10.1371/journal.pone.0051218

Lagrené, K., Bécu, M., Seiple, W., Raphanel, M., Combariza, S., Paques, M., Aubois, A., Duclos, B., Eandi, C., Girmens, J.-F., Mohand-Said, S., & Arleo, A. (2019). Healthy and pathological visual aging in a French follow-up cohort study. Investigative Opthalmology & Visual Science, 60, 5915–5915.

Lallart, E., Jouvent, R., Herrmann, F. R., Beauchet, O., & Allali, G. (2012). Gait and motor imagery of gait in early schizophrenia. Psychiatry Research, 198(3), 366–370. 10.1016/j.psychres.2011.12.013

Malouin, F., & Richards, C. L. (2010). Mental practice for relearning locomotor skills. In Physical Therapy (Vol. 90, Issue 2, pp. 240–251). 10.2522/ptj.20090029

Malouin, F., Richards, C. L., & Durand, A. (2010). Normal Aging and Motor Imagery Vividness: Implications for Mental Practice Training in Rehabilitation. Archives of Physical Medicine and Rehabilitation, 91(7), 1122–1127. 10.1016/j.apmr.2010.03.007

Marchetti, R., Vaugoyeau, M., Colé, P., & Assaiante, C. (2022). A sensorimotor representation impairment in dyslexic adults: A specific profile of comorbidity. Neuropsychologia, 165, 108134. 10.1016/j.neuropsychologia.2021.108134

Masters, R. S. W., Eves, F. F., & Maxwell, J. P. (2005). Development of a movement specific Reinvestment Scale. http://hub.hku.hk/handle/10722/115113

McAvinue, L. P., & Robertson, I. H. (2008). Measuring motor imagery ability: A review. European Journal of Cognitive Psychology, 20(2), 232–251. 10.1080/09541440701394624

Miall, R. C., & Wolpert, D. M. (1996). Forward Models for Physiological Motor Control. Neural Networks, 9(8), 1265–1279. 10.1016/S0893-6080(96)00035-4

Montero-Odasso, M., van der Velde, N., Martin, F. C., Petrovic, M., Tan, M. P., Ryg, J., Aguilar-Navarro, S., Alexander, N. B., Becker, C., Blain, H., Bourke, R., Cameron, I. D., Camicioli, R., Clemson, L., Close, J., Delbaere, K., Duan, L., Duque, G., Dyer, S. M., … the Task Force on Global Guidelines for Falls in Older Adults. (2022). World guidelines for falls prevention and management for older adults: A global initiative. Age and Ageing, 51(9), afac205. 10.1093/ageing/afac205

Nakano, H., Murata, S., Shiraiwa, K., & Nonaka, K. (2020). Increased time difference between imagined and physical walking in older adults at a high risk of falling. Brain Sciences, 10(6). 10.3390/brainsci10060332

Paraskevoudi, N., Balcı, F., & Vatakis, A. (2018). “Walking” through the sensory, cognitive, and temporal degradations of healthy aging. In Annals of the New York Academy of Sciences. 10.1111/nyas.13734

Personnier, P., Kubicki, A., Laroche, D., & Papaxanthis, C. (2010). Temporal features of imagined locomotion in normal aging. Neuroscience Letters, 476(3), 146–149. 10.1016/j.neulet.2010.04.017

Personnier, P., Paizis, C., Ballay, Y., & Papaxanthis, C. (2008). Mentally represented motor actions in normal aging: II. The influence of the gravito-inertial context on the duration of overt and covert arm movements. Behavioural Brain Research, 186(2), 273–283. 10.1016/J.BBR.2007.08.018

Podsiadlo, D. R., Richardson, S. (1991). The Timed Up and Go: A Test of Basic Functional Mobility for Frail Elderly Persons. Journal of the American Geriatrics Society, 39(2), 142–148.

Ren, J., Wu, Y. D., Chan, J. S. Y., & Yan, J. H. (2013). Cognitive aging affects motor performance and learning. Geriatrics & Gerontology International, 13(1), 19–27. 10.1111/j.1447-0594.2012.00914.x

Robin, N., Coudevylle, G. R., Dominique, L., Rulleau, T., Champagne, R., Guillot, A., & Toussaint, L. (2021). Translation and validation of the movement imagery questionnaire-3 second French version. Journal of Bodywork and Movement Therapies, 28, 540–546. 10.1016/j.jbmt.2021.09.004

Saimpont, A., Malouin, F., Tousignant, B., & Jackson, P. L. (2013). Motor imagery and aging. Journal of Motor Behavior, 45(1), 21–28. 10.1080/00222895.2012.740098

Saimpont, A., Malouin, F., Tousignant, B., & Jackson, P. L. (2015). Assessing motor imagery ability in younger and older adults by combining measures of vividness, controllability and timing of motor imagery. Brain Research, 1597, 196–209. 10.1016/j.brainres.2014.11.050

Sakurai, R., Fujiwara, Y., Yasunaga, M., Suzuki, H., Sakuma, N., Imanaka, K., & Montero-Odasso, M. (2017). Older adults with fear of falling show deficits in motor imagery of gait. *The Journal of Nutrition*, Health and Aging, 21(6), 721– 726. 10.1007/s12603-016-0811-1

Schott, N. (2012). Age-related differences in motor imagery: Working memory as a mediator. Experimental Aging Research, 38(5), 559–583. 10.1080/0361073X.2012.726045

Schott, N., & Munzert, J. (2007). Temporal accuracy of motor imagery in older women. International Journal of Sport Psychology, 38(3), 304–320.

Seidler, R. D., Bernard, J. A., Burutolu, T. B., Fling, B. W., Gordon, M. T., Gwin, J. T., Kwak, Y., & Lipps, D. B. (2010). Motor control and aging: Links to age-related brain structural, functional, and biochemical effects. Neuroscience & Biobehavioral Reviews, 34(5), 721–733. 10.1016/j.neubiorev.2009.10.005

Shadmehr, R., & Krakauer, J. W. (2008). A computational neuroanatomy for motor control. Experimental Brain Research, 185(3), 359–381. 10.1007/s00221-008-1280-5

Skoura, X., Papaxanthis, C., Vinter, A., & Pozzo, T. (2005a). Mentally represented motor actions in normal aging: I. Age effects on the temporal features of overt and covert execution of actions. Behavioural Brain Research, 165(2), 229–239. 10.1016/j.bbr.2005.07.023

Snyder, B. A., Munter, A. D., Houston, M. N., Hoch, J. M., & Hoch, M. C. (2016). Interrater and intrarater reliability of the semmes-weinstein monofilament 4-2-1 stepping algorithm: Reliability of the 4-2-1 Algorithm. Muscle & Nerve, 53(6), 918–924. 10.1002/mus.24944

Stevens, J. A., Mack, K. A., Paulozzi, L. J., & Ballesteros, M. F. (2008). Self-Reported Falls and Fall-Related Injuries Among Persons Aged ≥65 Years–United States, 2006. Journal of Safety Research, 39(3), 345–349. 10.1016/j.jsr.2008.05.002

Subirats, L., Allali, G., Briansoulet, M., Salle, J. Y., & Perrochon, A. (2018). Age and gender differences in motor imagery. Journal of the Neurological Sciences, 391, 114–117. 10.1016/j.jns.2018.06.015

Vandevoorde, K., & De Xivry, J.-J. O. (2020). Does somatosensory acuity influence the extent of internal model recalibration in young and older adults? Neuroscience. 10.1101/2020.10.16.342295

Watanabe, M., & Tani, H. (2022). Using crutches during walking possibly reduces gait imagery accuracy among healthy young and older adults. Journal of Physical Therapy Science, 34(10), 673–677. 10.1589/jpts.34.673

Williams, S., Guillot, A., Di Rienzo, F., & Cumming, J. (2015). Comparing self-report and mental chronometry measures of motor imagery ability. European Journal of Sport Science, 15, 1–9. 10.1080/17461391.2015.1051133

Wolpert, D. M., Diedrichsen, J., & Flanagan, J. R. (2011). Principles of sensorimotor learning. Nature Reviews Neuroscience, 12(12), 739–751. 10.1038/nrn3112

Wolpert, D. M., Ghahramani, Z., & Jordan, M. I. (1995). An Internal Model for Sensorimotor Integration. Science, 269(5232), 1880–1882. 10.1126/science.7569931

Zabicki, A., de Haas, B., Zentgraf, K., Stark, R., Munzert, J., & Krüger, B. (2019). Subjective vividness of motor imagery has a neural signature in human premotor and parietal cortex. NeuroImage, 197, 273–283. 10.1016/j.neuroimage.2019.04.073

Zapparoli, L., Mariano, M., & Paulesu, E. (2022). How the motor system copes with aging: A quantitative meta-analysis of the effect of aging on motor function control. Communications Biology, 5(1), Article 1. 10.1038/s42003-022-03027-2

Zhuang, X., Zhang, T., Chen, W., Jiang, R., & Ma, G. (2020). Pedestrian estimation of their crossing time on multi-lane roads. Accident Analysis & Prevention, 143, 105581. 10.1016/j.aap.2020.105581

